# Improved daylight vision following AAV-mediated expression of *R9AP* in murine rod photoreceptors

**DOI:** 10.1101/2020.02.19.955765

**Authors:** Koji M. Nishiguchi, Kosuke Fujita, Enrico Cristante, James W. Bainbridge, Ronald H. Douglas, Toru Nakazawa, Alexander J. Smith, Robin R. Ali

**Author notes:** Corresponding author Robin R. Ali, Department of Genetics, University College London Institute of Ophthalmology, 11-43 Bath Street, London, EC1V 9EL, UK.

## Abstract

Cone photoreceptors mediate daylight vision and are the primary cells responsible for vision in humans. Cone dysfunction leads to poor quality daylight vision because rod photoreceptors become saturated and non-functional at high light levels. Here we demonstrate that in mice lacking cone function, AAV-mediated over-expression of Rgs9-anchor protein (R9AP), a critical component of the GTPase complex that mediates the deactivation of the phototransduction cascade, results in desensitization of rod function and a “photopic shift” of the rod-driven electroretinogram. This treatment enables rods to respond to brighter light (up to ∼2.0 log) with increased visually-evoked cortical responses to high intensity stimulation. These results suggest that AAV-mediated transfer of *R9ap* into rods might be used to improve daylight vision in humans visually handicapped by cone dysfunction.

## Introduction

In most mammals, including mice and humans, rod photoreceptors, which mediate vision in dim light, far outnumber cone photoreceptors ^1^. However, for humans in an industrialized world where artificial illumination facilitates cone function throughout the day, rod-mediated vision is less critical. Many individuals with congenitally absent rod function are identified only incidentally and, in fact, may not recognize their abnormal vision^2, 3^. In contrast, individuals affected by cone dysfunction are typically symptomatic and often suffer visual impairment dependent on the degree of their cone dysfunction. In some conditions, when the cones are lost or dysfunctional, rods remain well preserved. Although rods are able to detect very low levels of light, they become saturated at high light levels. Cones, on the other hand, are less sensitive, but are capable of processing large amounts of light and are continuously functional in bright daylight. This difference is, in part, due to the efficiency of the deactivation machinery of phototransduction, the GTPase complex composed of RGS9, R9AP (also known as RGS9BP), and Gß5^4, 5^. RGS9 is the catalytic component that hydrolyses the GTP coupled to the G-protein, whereas R9AP and Gß5 are the essential constitutive subunits^4, 5^. Importantly, R9AP tethers the complex to the disc membrane in the photoreceptor outer segment where phototransduction signaling takes place ^6^. Expression of R9AP determines the level of the GTPase complex, such that any RGS9 produced in excess of R9AP is quickly degraded ^7^. Over-expression of R9AP in murine rods is sufficient to increase the GTPase activity and to substantially increase the speed of their deactivation kinetics as evidenced by single cell recordings^8^ and flicker electroretinogram (ERG) responses in transgenic mice ^9^. In cones, RGS9 expression has been estimated to be ∼10-fold higher than in rods^10, 11^. This provides a basis for the ability of the cones to recover quickly from light exposure and thus maintain function in response to a continuous light stimulus. It also allows cones to respond to more rapid stimulation. Indeed, patients with bradyopsia, who have delayed deactivation of the phototransduction cascade caused by genetic defects in RGS9 or R9AP, have a profound impairment of cone-mediated vision including day blindness and reduced ability to see moving objects^12, 13^. Rod-mediated vision is less affected by the same mutation. Cone dysfunction/degeneration with relatively preserved rods are observed in a few forms of retinal dystrophies, such as cone dystrophy, complete and incomplete achromatopsia, and cone-rod dystrophy ^14, 15, 16, 17, 18^, as well as more frequent disease such as early-onset high myopia^19, 20^. In these conditions, patients are often obliged to rely heavily on rod mediated vision even under light resulting in from debilitating photophobia of unknown cause and day blindness caused by low bleaching threshold of the rods.

In this study, we sought to define the therapeutic potential of AAV-mediated transfer of the murine *R9ap* gene into rods using murine models lacking cone vision. Electrophysiological and behavioral tests showed that vector-mediated R9AP over-expression in adult mice can increase the refresh rate of rods, allowing them to respond to higher light levels thereby improving daylight vision.

## Results

### R9AP over-expression in rods and increased speed of photoreceptor deactivation

To study the effect of AAV-mediated R9AP over-expression on the GTPase complex in the rods, the level and distribution of the catalytic component, RGS9, was examined following subretinal injection of rAAV2/8.Rho.mR9ap in *Cnga3-/-* mice. These mice have normal rod function but absent cone function and serve as an animal model of achromatopsia, a rare but most severe form of cone dystrophy with a complete absence of cone function from birth. Four weeks later, the retina showed increased immunoreactivity against RGS9 in treated compared to untreated eyes (Fig. 1A). This was recognized most obviously as a difference in the distribution of the RGS9. In the treated eyes, RGS9 expression was recognized throughout the entire photoreceptor layer, including the outer nuclear layer and outer plexiform layer, whereas the expression was confined to the inner and the outer segments in the untreated contralateral eye. Western blot analysis confirmed the increased RGS9 protein expression in the treated retina (Fig. 1B). These results confirm the prediction that an over-expression of R9AP increases the level of RGS9 and the GTPase complex.

**Figure 1.**
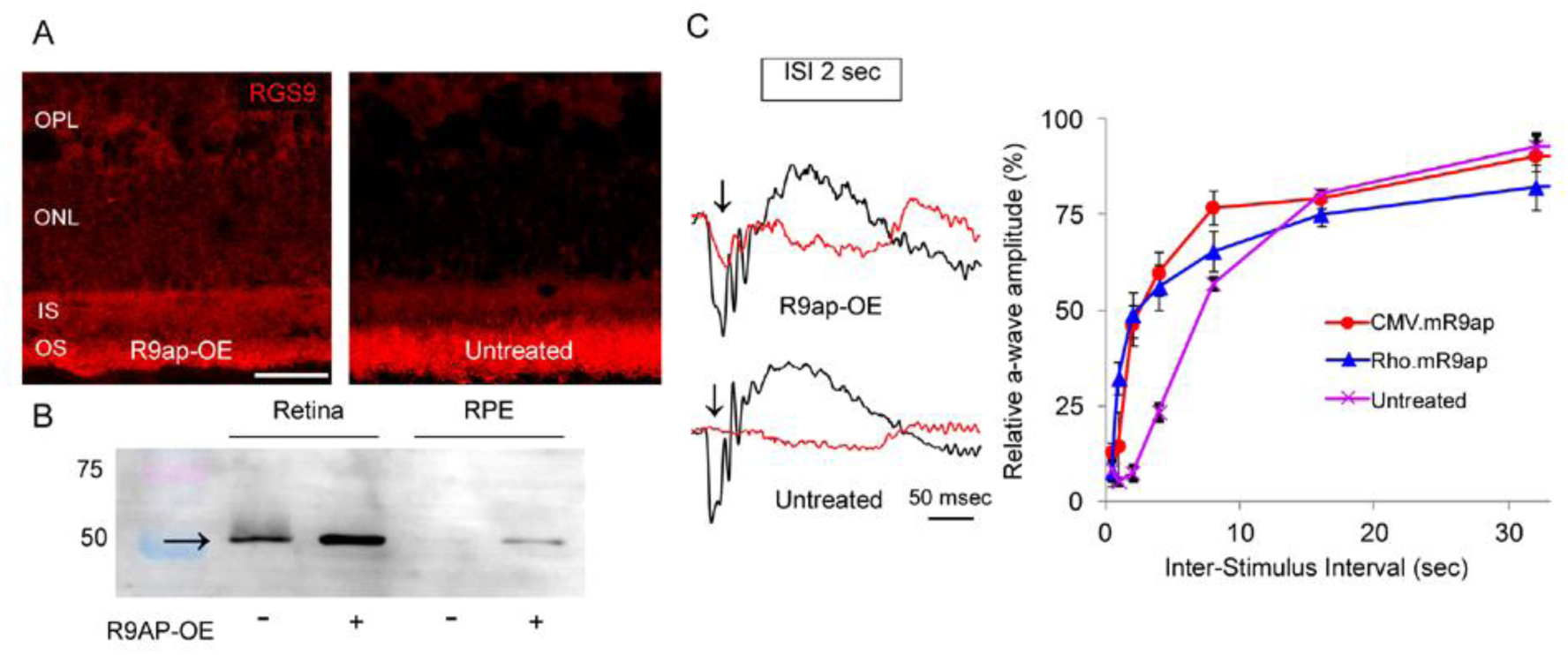
AAV-mediated R9AP over-expression in rods and accelerated a-wave deactivation. **A**. Increased RGS9 expression in a *Cnga3-/-* eye treated with rRAAV2/8.Rho.mR9ap. Over-expression of R9AP results in increased immunoreactivity toward RGS9 (red) throughout the photoreceptor layer in the treated eye (left) compared to the untreated (right) under light adaptation. This was recognized most obviously as a difference in the distribution of the RGS9. In the treated eyes, RGS9 expression was recognized throughout the entire photoreceptor layer, including the outer nuclear layer and outer plexiform layer, whereas the expression was confined to the inner and the outer segments in the untreated contralateral eye. **B**. Westernblot shows increased expression of RGS9 in the retina of an eye over-expressing R9AP (bottom). A small amount of RGS9 protein was also detected in the RPE of the treated eye. This may reflect “spill over” of the excessive protein contained within the phagocytosed disc membrane. “+” indicates the eyes over-expressing R9AP and “-” indicates the untreated contralateral eye. Scale bar indicates 25 µm. **C**. Increased speed of ERG a-wave amplitude recovery in the *Cnga3-/-* eyes treated with rAAV2/8.Rho.mR9ap and rAAV2/8.CMV.mR9ap. Representative responses to a probe flash (black traces) and a 2nd flash (red traces) with an inter-stimulus interval (ISI) of 2 seconds from the treated (top) and untreated (bottom) eyes from the same animal. Note that a second flash yields a clearly visible small a-wave (arrow) in the treated eye, whereas an a-wave is not apparent (arrow) in the untreated fellow eye. A plot of a-wave recovery at various ISIs in the treated and untreated eyes is also shown. The eyes injected with rAAV2/8.CMV.mR9ap (n = 5) or rAAV2/8.Rho.mR9ap (n = 7) have faster recovery kinetics than untreated eyes (n = 5) which is most apparent at shorter ISIs. The data is presented as average ± standard error of the mean. OE: over-expression.

Next we studied the functional effect of R9AP over-expression on rod phototransduction using paired-flash ERGs^21^. In this paradigm, a pair of flashes of equal intensity are delivered with a variable inter-stimulus interval and the recovery of the second response relative to the first is measured. In the rod photoreceptors, the speed of the a-wave (originating from photoreceptors) recovery is dependent on the deactivation speed of their phototransduction signaling^21^. The time constant (σ) for 50% recovery of a-wave amplitude was reduced by ∼60% in the *Cnga3-/-* eyes injected with rAAV2/8.CMV.mR9ap (σ = ∼ 2.99 sec) compared to the untreated eyes (σ = ∼ 7.38 sec; Fig. 1C). Similarly, a reduced a-wave recovery time was observed when a rhodopsin promoter was used to drive over-expression of R9AP (rAAV2/8.Rho.mR9ap; σ = ∼ 2.74 sec; Fig. 1C) in the same mouse line (*Cnga3-/-*) or the same virus (rAAV2/8.CMV.mR9ap) in another cone-defective mouse line (*Pde6c-/-*; Supplementary Fig. S1). These observations indicate that the AAV-mediated over-expression of R9AP can significantly increase the deactivation speed of rod phototransduction, thus shortening their photoresponse through increasing the level of RGS9 and the GTPase complex.

### Photopic shift of rod function by over-expression of R9AP

To define the effects of an increased GTPase activity and deactivation speed of phototransduction achieved through the overexpression of R9AP on the rod photoreceptor function, dark-adapted 6 Hz flicker ERGs were recorded using flashes of incremental intensities. In this paradigm, untreated rods respond to flashes within a defined range of light intensities and do not show measurable light responses to the brightest lights where their recovery speed cannot catch up with the incoming photons (Fig. 2A). Therefore, the test provides an objective index of the operating range of the rods. The eyes over-expressing R9AP through injection of rAAV2/8.CMV.mR9ap or rAAV2/8.Rho.mR9ap showed increased responses to brighter flashes compared to untreated contralateral eyes. This was accompanied by an elevation of the upper threshold of the rod response by up to ∼2 log units (Fig. 2A), with little effect on the maximal photoresponse (151 ± 17 µV in the treated vs 162 ± 29 µV in the untreated eyes; average ± standard error of the mean). This “photopic shift” in the operating range of the rods was coupled with a reciprocal elevation of the lower threshold of the response by up to ∼1.5 log units. We obtained similar results when R9AP was over-expressed in *Pde6c-/-* mice (Supplementary Fig. S2). These fundamental changes in operating range following the treatment allowed the rods to respond to flashes of longer durations (Fig. 2B) and to flashes under a constant white background illumination (30 cd/m^2^) used to saturate normal rod function completely and isolate cone function in a standard ERG protocol (Fig. 2C). Taken together, these results establish that R9AP-over expression in rods results in their desensitization and allows them to mediate photopic function under conditions where normal rods are virtually non-functional in exchange for the loss of scotopic function. Meanwhile, the treatment of wildtype mice using the same viral vectors failed to show a measurable change in retinal function (Supplementary Fig. S3).

**Figure 2.**
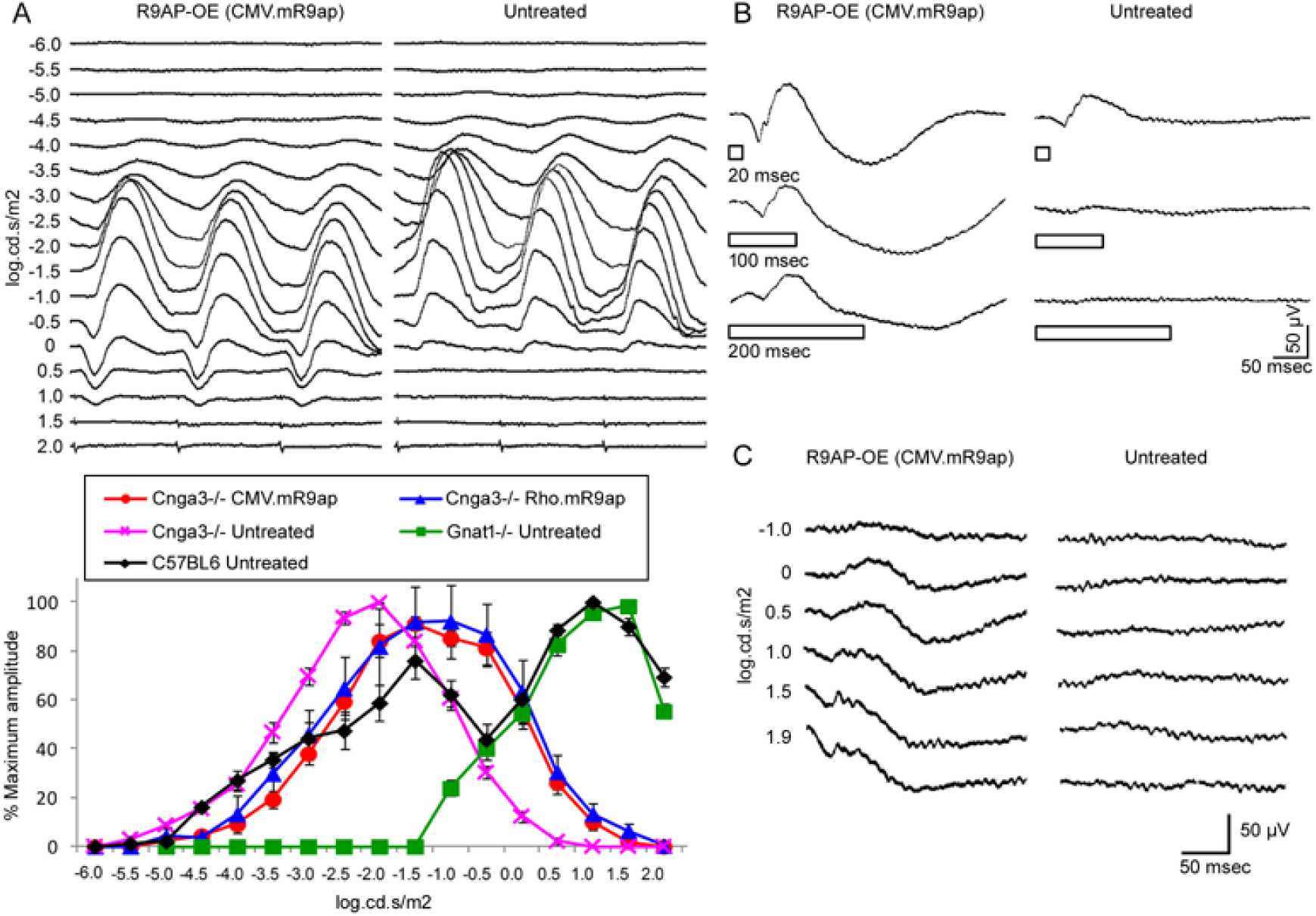
Gain of photopic function by rods through over-expression of R9AP. **A**. Elevation of response threshold and “photopic shift” of 6Hz ERGs through over-expression of R9AP in rods of *Cnga3-/-* mice. Representative 6Hz ERG traces from a *Cnga3-/-* mouse in which one eye was treated with rAAV2/8.CMV.mR9ap and the other eye was left untreated (top panel). ERG traces are aligned from responses against the dimmest flash (−6.0 log cd.s/m^2^) to the brightest flash (2.0 log cd.s/m^2^; bottom) from the top to the bottom in 0.5 log.cd.s/m^2^ steps. Note that the lower threshold flash intensity at which the responses emerge is increased, which is coupled with an elevated response threshold to brighter flashes. This results in a “photopic shift” of retinal function in the eye treated with rAAV2/8.CMV.mR9ap. The bottom panel shows a summary of 6Hz ERG results demonstrating a photopic shift in retinal function following treatment with rAAV2/8.CMV.mR9ap or rAAV2/8.Rho.mR9ap. ERG responses from *Gnat1-/-* mice deficient in rod function show cone-mediated function. Meanwhile, responses from C57BL6 mice are derived from both rod and cone photoreceptors. The data is presented as % amplitude relative to the maximal response and is shown as average ± standard error of the mean. *Cnga3-/-* eyes treated with rAAV2/8.CMV.mR9ap (Cnga3-/- CMV.R9ap; N = 8), *Cnga3-/-* eyes treated with rAAV2/8.Rho.mR9ap (Cnga3-/- Rho.R9ap; N = 6), untreated *Cnga3-/-* eyes (Cnga3-/- Untreated; N = 8), untreated *Gnat1-/-* eyes (Gnat1-/- Untreated; N = 6), and untreated wildtype eyes (C57BL6 Untreated; N = 6). **B**. Increased retinal responses to long flashes in an *Cnga3-/-* eye treated with rAAV2/8.CMV.mR9ap. Open rectangles denote the duration of the flash. Note that in the eye treated with rAAV2/8.CMV.mR9ap, responses are detectable with increased durations of the light stimulus. Conversely, the untreated contralateral eye shows little or no response when recorded simultaneously under identical conditions. **C**. Gain of retinal function under photopic conditions in *Cnga3-/-* eyes treated with rAAV2/8.CMV.mR9ap. Note that the treated rods show responses under photopic conditions (white background light of 30 cd/m^2^), whereas the untreated rods in the contralateral eye of the same mouse recorded simultaneously remain unresponsive.

### Delayed and reduced transmission of the altered rod response by rod bipolar cells

An overexpression of R9AP in rods results in faster photoreceptor deactivation kinetics, which allows them to respond to a larger amount of photons. Meanwhile, the accelerated deactivation should also result in a shorter duration of neurotransmitter release at the rod spherule, which may in turn reduce the transmission efficiency of the neural signal to the downstream bipolar cells. To this end, we studied the speed and magnitude of signal transmission from photoreceptors to bipolar cells by analyzing implicit time and amplitudes of the ERG a-wave (originating from the photoreceptors) and the b-wave (originating from bipolar cells) elicited by a short isolated flash in *Cnga3-/-* mice treated with rAAV2/8.Rho.mR9ap (Fig. 3). The a-wave trough marks the point at which the bipolar cell-driven b-wave becomes detectable. Some increase in the b-wave implicit times without notable change in a-wave implicit times, most likely reflecting a reduced transmission efficiency of the visual signal at the level of the rod bipolar cells, was observed. In addition, b-wave amplitudes probing bipolar cell responses were also reduced. This indicates that the amount of signal transmission to a flash of a given intensity is reduced in treated animals. However, as there is a relatively large variation of ERG responses between normal individuals, a small delay or reduction in rod response does not necessarily translate into noticeable visual dysfunction ^22^.

**Figure 3.**
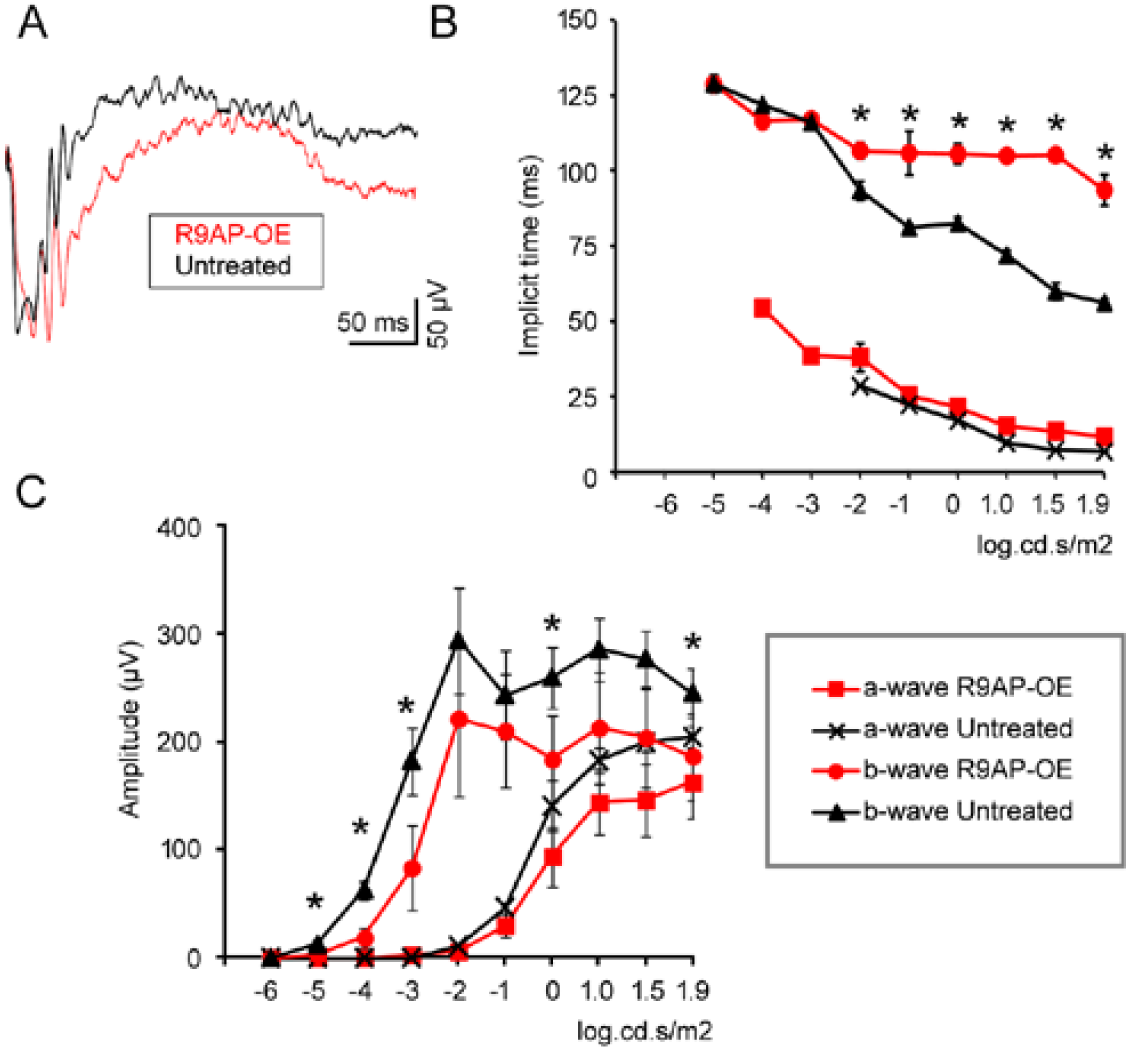
Efficient transmission of the altered photoreceptor signal to rod bipolar cells in eyes over-expressing R9AP. **A**. Representative ERG traces recorded after rAAV2/8.CMV.mR9ap injection (red trace) in a *Cnga3-/-* mouse using a saturating flash (1.9 log cd.s/m^2^). The contralateral eye served as an untreated control (black trace). **B**. Delayed b-wave peaks in eyes over-expressing R9AP. A-wave and b-wave implicit times were measured from ERG responses in the treated (red symbols) and the untreated (black symbols) *Cnga3-/-* eyes (N=5 each). B-wave peak was significantly delayed in response to flashes of higher intensities whereas no difference was observed in a-wave implicit times. **C**. Reduced b-wave amplitudes recorded from *Cnga3-/-* mice (N=5) in eyes over-m expressing R9AP. A-wave and b-wave amplitudes were measured from ERG response in the treated (red symbols) and the untreated (black symbols) *Cnga3-/-* eyes (N=5 each). Note b-wave amplitudes were significantly reduced in response to flashes of various intensities. All data are presented as average ± standard error of the mean. OE: over-expression. * P < 0.05.

### Photopic shift of rod-mediated cortical responses by over-expression of R9AP

An overexpression of R9AP in rods results in desensitization of the cells and a “photopic shift” of their function. This is accompanied by a small delay and reduction in the bipolar cell response, which may or may not affect visual perception. In order to assess the functional consequence of R9AP overexpression in rods at the level of the visual cortex, visually evoked potentials (VEP) were recorded. *Pde6c-/-* mice treated with rAAV2/8.Rho.mR9ap were dark-adapted and presented with series of flashes of incremental intensities. The intensity-response profile showed increased P1-N1 and N1-P2 amplitudes particularly at higher flash intensities, consistent with a “photopic shift” of rod-mediated cortical responses by the over-expression of R9AP (Fig. 4). Interestingly, the magnitude of the cortical responses as measured by the amplitudes were not reduced at all flash intensities, indicating that the delay and reduction of bipolar responses observed using ERG have minimal effects on the visual perception mediated by altered rod function at some intensities.

**Figure 4.**
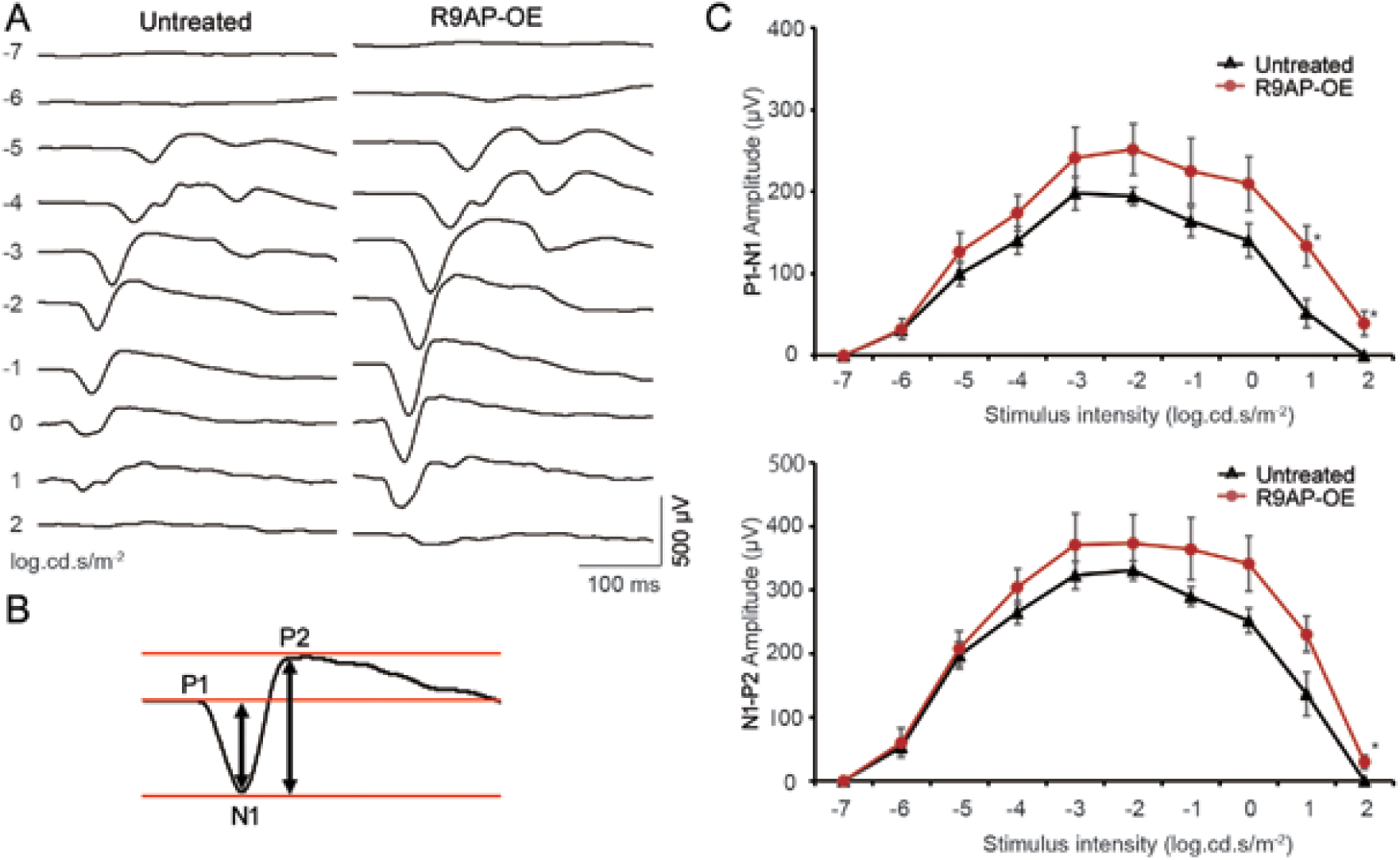
Increased cortical responses to bright flashes following R9AP over-expression in rods. **A**. Representative VEPs in response to flashes of various intensities from a *Pde6c-/-* mouse in which one eye was treated with rAAV2/8.CMV.mR9ap and the other eye was left untreated. **B**. Schematic illustration of P1-N1 and N1-P2 components of a VEP trace and measurement of their amplitudes. **C**. Increased amplitudes of P1-N1 (top) and N1-P2 (bottom) components following treatment with rAAV2/8.CMV.mR9ap at high stimulus intensities. N = 5. Data with error bars were presented as mean ± standard error of the mean. * P < 0.05.

## Discussion

In the absence of functional cone photoreceptors, vision is dependent on the rods, which normally only function efficiently in lower light conditions. Hence, in individuals with a substantial loss of cones, the low bleaching threshold of remaining rods severely limits the quality of daylight vision. The principle idea of the current study is to “shift” the functional range of the rods towards that of cones through viral vector-mediated over-expression of R9AP, with the aim of compensating for cone dysfunction. The key advantage of this therapeutic approach lies in the fact that the sensitivity of rods is altered through enhancement of a well-defined endogenous regulatory mechanism. This also means that the treated rods may efficiently utilize the downstream rod neural circuit in the retina and in the brain, which ensures the visual system can accommodate the altered retinal function. Importantly, a recent study has indicated unexpectedly well preserved plasticity of rod-mediated visual pathway by showing full restoration of rod-mediated visual acuity in blind adult mice of up to 9 months of age ^23^. The treatment relies on the functional augmentation of the abundant rods and does not require the presence of cones. Therefore, therapeutic application may be considered when extensive dysfunction exists due to substantial loss of cones. Thus, forms of cone dystrophy/achromatopsia, cone-rod dystrophy, and acquired cone dysfunction ranging from high myopia to diabetic macular edema may be the suitable targets, as a progressive loss of the cones is documented in patients with these conditions ^14, 15, 19, 20^. However, older achromatopsia patients may also benefit from the rod augmentation approach described herein. Not only has some progressive loss of cones been described in achromatopsia patients ^15, 24, 25^, but the lack of cone input from birth likely affects the development of physiological cone-dependent visual neuronal circuits including those of the visual cortex^26^, limiting the potential for older individuals to benefit from therapeutic restoration of retinal function.

The obvious drawback of the current approach is the inability to restore high resolution vision in humans which is dependent on the utilization of densely packed cones driving post-receptoral neural circuits in the rod-free fovea. An additional drawback is the persistence of colour vision defects, restoration of which depends on the use of cone visual pigments of different absorption spectra. As the “shift” of rod function involves a desensitization of up to ∼2.0 log units, patients may be able to see objects in environment up to ∼100 times brighter than before. However, the operating range of intact cones appears to be at least another ∼2.0 log units higher than the R9AP-overexpressing rods (Fig. 2). Thus, the current approach is only partially effective at compensating for the absence of cones.

In conclusion, our approach demonstrates that AAV-mediated over-expression of R9AP in rod photoreceptors might be an effective therapeutic approach to ameliorate cone dysfunction, reducing hypersensitivity to light and improving daylight vision in people affected by advanced cone degeneration caused, for example, by conditions such as inherited disease or wide range of acquired macular dysfunction and degeneration in which cone-mediated vision is severely affected.

## Methods

### Animals

C57BL6 (Harlan, UK), *Cnga3*^*cpfl5/cpfl5*^ (referred herein as *Cnga3*^*-/*^; J.R. Heckenlively, University of Michigan), *Pde6c*^*cpfl1/cpfl1*^ (referred herein as *Pde6c*^*-/-*^; J.R. Heckenlively, University of Michigan, MI) ^27^, and *Gnat1*^*-/-*^ (J. Lem, Tufts University School of Medicine, MA) ^28^ mice were maintained in the animal facility at University College London. Mice lines, C57BL6 (SLC, Shizuoka, Japan), *Cnga3*^*cpfl5/cpfl5*^ (Jackson Laboratory, ME), and *Pde6c*^*cpfl1/cpfl1*^ (Jackson Laboratory), were separately obtained for the experiments related to measurements of visually-evoked potentials. Adult male and female animals were 6-12 weeks old at the time of viral injection and were used for experiments at least 2 weeks after the injection to allow for a sufficient expression of R9AP. All the mice used were between ages of 2 to 6 months and were age matched between groups for a given experiment. All experiments were conducted in accordance with the Policies on the Use of Animals and Humans in Neuroscience Research and with the ARVO Statement for the Use of Animals in Ophthalmic and Vision Research. Animals were kept on a standard 12/12 hour light-dark cycle.

### Plasmid constructions and production of recombinant AAV8

The murine *R9ap* cDNA was PCR amplified from murine retinal cDNA using primers designed to encompass the whole of the coding region. The *R9ap* cDNA was cloned between the promoter (CMV promoter or bovine rhodopsin promoter) and the SV40 polyadenylation site. These plasmids were used to generate two pseudotyped AAV2/8 viral vectors, rAAV2/8.CMV.mR9ap and rAAV2/8.Rho.mR9ap, as described previously ^29^.

In brief, recombinant AAV2/8 vector was produced through a triple transient transfection method as described previously^30^. The plasmid construct, AAV serotype-specific packaging plasmid and helper plasmid were mixed with polyethylenimine to form transfection complexes which was then added to 293T cells and left for 72 h. The cells were harvested, concentrated and lysed to release the vector. The AAV2/8 was purified by affinity chromatography and concentrated using ultrafiltration columns (Sartorius Stedim Biotech, Goettingen, Germany), washed in PBS and concentrated to a volume of 100–150 µl. Viral particle titres were determined by dot-blot or by real-time PCR. Purified vector concentrations used were 1-2 × 10^12^ viral particles/ml.

### Immunohistochemistry

Six weeks after unilateral subretinal injection of rAAV2/8.Rho.mR9ap, both eyes from a *Cnga3-/-* mouse were quickly removed and snap frozen in liquid nitrogen. After cryoembedding the eye in OCT (RA Lamb, Eastborne, UK), the eyes were cut as 15μm transverse sections and air-dried for 15 - 30 min. For immunohistochemistry, sections were pre-blocked in PBS containing normal donkey serum (2%), bovine serum albumin (2 %) 1 hr before being incubated with anti-RGS9 antibody (1:500; Santa Cruz Biotechnology, SantaCruz, CA) for 2 hours at room temperature. After rinsing 2 × 15 min with PBS, sections were incubated with the appropriate Alexa 546-tagged secondary antibody (Invitrogen, Carlsbad, CA) for 2 hrs at room temperature (RT), rinsed and counter-stained with Hoechst 33342 (Sigma-Aldrich, Gillingham, UK). Retinal sections were viewed on a confocal microscope (Leica TCS SP2, Leica Microsystems; Wetzlar, Germany).

### Western blotting

The eyes from a 4-week old *Cnga3-/-* mouse after unilateral sub-retinal injection of rAAV2/8.Rho.mR9ap were collected. After separating the neural retina from the RPE/choroid/sclera complex, tissues were homogenized in RIPA buffer (5 µl/mg tissue) with protease inhibitors (Sigma Aldrich, UK), agitated at 4°C for 30 mins and subsequently centrifuged at 17000xg for 30 mins at 4°C. Supernatants were stored at −80°C until further use. Protein species were resolved as previously described ^31^. Briefly, equal amounts of protein extracts (10 μg, denatured in Laemmli’s loading buffer) were loaded and resolved on a two-layer SDS-PAGE gel (12% resolving gel; 4% loading gel) together with pre-stained standards (ThermoFisher Scientific, UK). Separated proteins were electro-transferred to PVDF membranes (Millipore, UK) for 1 hour. Membranes were blocked for 1 hr at RT in 3% BSA PBS 0.1% Tween-20 (PBS-T) and then incubated overnight at 4°C with primary antibody (rabbit anti-RGS9, 1:500 in blocking solution; Abcam, UK). After washing three times in PBS-T, membranes were incubated in secondary antibody for 1 hr at RT (goat anti-rabbit HRP conjugated, 1:2500 in blocking solution; ThermoFisher Scientific, UK).

Immune-reactive bands were visualized by enhanced chemiluminescence (ECL plus GE Healthcare, UK). A Fujifilm LAS-1000 Luminescence Image Analyser was used to detect the signal after 1 minute incubation in the reaction mix, by exposing the membranes for 1 – 60 seconds.

### Electroretinogram (ERG)

ERGs were recorded from both eyes after mice were dark adapted overnight using a commercially available system (Espion E2, Diagnosys LLC, Lowell, MA) as described previously^32^. The animals were anesthetized with an intraperitoneal injection of a 0.007 ml/g mixture of medetomidine hydrochloride (1 mg/ml), ketamine (100 mg/ml), and water at a ratio of 5:3:42 before recording. Pupils were fully dilated using 2.5% phenylephrine and 1.0% tropicamide. Midline subdermal ground and mouth reference electrodes were first placed, followed by positive silver electrodes that were allowed to lightly touch the center of the corneas under dim red illumination. A drop of Viscotears 0.2% liquid gel (Dr. Robert Winzer Pharma/OPD Laboratories, Watford, UK) was placed on top of the positive electrodes to keep the corneas moistened during recordings and the mouse was allowed to further dark-adapt for 5 minutes. Bandpass filter cutoff frequencies were 0.312 Hz and 1000 Hz. Recovery speed of photoresponse was measured using paired flash paradigm where pairs of flashes with identical saturating intensity (1.8 log.cd.s/m^2^) separated by various inter-stimulus intervals (ISI; 0.5, 1, 2, 4, 8, 16, 32, 64 sec) were presented. In this paradigm, the 1^st^ flash would completely suppress the electric responses of rod mechanisms which allow observation of the speed of functional recovery of the rod function by presenting 2^nd^ flash with different ISI. Sufficient amount of time (150 sec) were provided between pairs of flashes to allow full recovery of the 1^st^ flash. Then the recovery of a-wave amplitude observed should reflect the speed of deactivation of the rods in animals devoid of cone function since the flash should only bleach a fraction (0.02%) of the rhodopsin ^33, 34^. Scotopic 6 Hz flicker intensity series were performed as previously reported with a few modifications.^35^ We used 17 steps of flash intensities ranging from −6 to 2 log.cd.s/m^2^ each separated by 0.5 log unit. For each step, after 10 seconds of adaptation, 600 msec sweeps were averaged 20 times using the same flash condition. Series of dark-adapted responses were also obtained using longer flashes with durations of 20, 100, and 200 msec all at 83.3 cd/m^2^. Standard single flash scotopic recordings were obtained from dark-adapted animals at the following increasing light intensities: −6, −5, −4, −3, −2, −1, 0, 1.0, 1.5, and 1.9 log.cd.s/m^2^. Photopic flash recordings were performed following 5 min light adaptation intervals on a background light intensity of 20 cd/m^2^, which was also used as the background light for the duration of the recordings. Photopic light intensities used were −2, −1, 0, 1, 1.5, and 1.9 log.cd.s/m^2^.

### Visually evoked potentials (VEP)

Before recording VEPs from the visual cortex, electrodes were surgically placed according to the following procedures^36^. Mice were anesthetized by a single intraperitoneal injection of medetomidine hydrochloride (0.6 mg/kg, Meiji Co. Ltd., Tokyo, Japan) and ketamine (36mg/kg, Daiichi-Sankyo, Tokyo, Japan). The part of scalp proximal to the bregma was shaved and disinfected by povidone iodine (Meiji Co. Ltd.). After excising this part of the skin, connective tissues were removed from bone using a scalpel blade. The recording electrodes were placed in the skull overlying the right and left primary visual cortex (3.6-mm caudal to bregma and 2.3-mm lateral) and reference electrode was fixed in the prefrontal cortex (2.0-mm rostral to bregma) ^37^. Before inserting screw electrodes, small holes (0.5mm diameter) were drilled onto the skull. Three stainless steel pan-head screws (M0.6 × 3mm length) were used for electrodes, which were screwed 1mm into the skull so that the tip of the screw electrode was in contact with the brain surface. These screws are then fixed with cyanoacrylate adhesives (Toa Gosei Co. Ltd., Tokyo, Japan). The ground electrode was clipped onto the tail. Mouse was kept warm on a heating pad through the procedure. After surgical procedures, mice were given a reversal agent, Atipamezole (0.35mg/kg, Meiji Co. Ltd.) and each animal was kept individually in a separate cage.

The VEP was recorded 7 days after surgically implanting the electrodes. First, mice were dark-adapted overnight, after which pupils were fully dilated with 2.5% phenylephrine and 1.0% tropicamide eye drops. The screw electrodes were connected with recording unit (PuREC, Mayo, Inazawa, Japan) synchronized with LED stimulation coupled with a Ganzfeld dome (LS-100, Mayo, Inazawa, Japan). White flash was presented with intensity increased at 1 log.cd.s/m^2^ step (range: −7 to 2 log.cd.s/m^2^). A band-pass filter (0.3Hz to 50Hz) was applied to the data upon acquisition of the electric responses.

## Acknowledgements

We thank Professor J.R. Heckenlively (University of Michigan) for the kind gift of *Cnga3*^*cpfl5/cpfl5*^ mice and *Pde6c*^*cpfl1/cpfl1*^ and Professor J. Lem (Tufts University School of Medicine) for *Gnat1-/-* mice. This work was supported by UK The Medical Research Council and RP Fighting Blindness, the NIHR Biomedical Research Centre at Moorfields Eye Hospital and UCL Institute of Ophthalmology, Takeda Science Foundation, and Novartis.

## Author contributions

K.M.N. F.K. E.C., R.H.D. and J.W.B.B. contributed to experimental execution. K.M.N., T.N. A.J.S and R.R.A. contributed to the concept and design of the experiments, funding and to manuscript writing.

