## Supplemental file for "Improved daylight vision following AAV-mediated expression of *R9AP* in murine rod photoreceptors"

### Supplementary Information

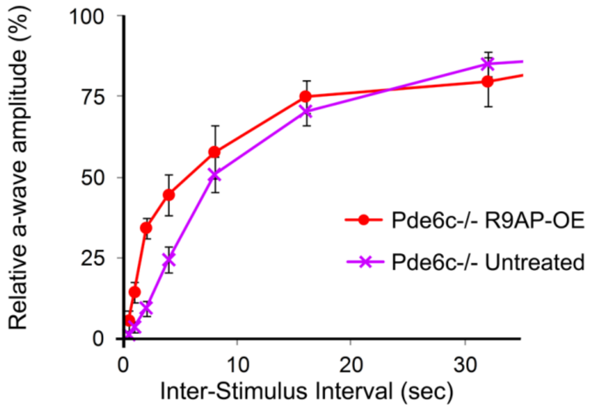


**Figure S1. Accelerated recovery of rod photoresponses by over-expression of R9AP in *Pde6c-/-* mice**

The time constant (σ) for 50% recovery of a-wave amplitude was reduced by ~50% in *Pde6c-/-* eyes injected with rAAV2/8.CMV.mR9ap (σ = ~ 5.75 sec) compared to untreated contralateral eyes (σ = ~ 11.46 sec), consistent with accelerated deactivation of phototransduction following the treatment. N = 6. Data with error bars were displayed as mean ± standard error of the mean.

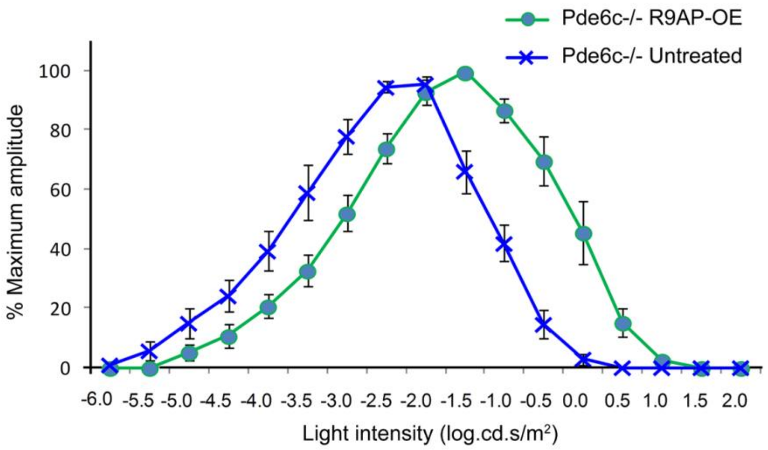


**Figure S2. "Photopic shift" of the intensity-response curve following over-expression of R9AP in *Pde6c-/-* mice**

The eyes injected with rAAV2/8.CMV.mR9ap showed a photopic shift of 6 Hz ERG responses to incremental flash intensities compared to untreated contralateral eyes in *Pde6c-/-* mice (N = 6). The data is presented as % amplitude relative to maximal response and is displayed as average ± standard error of the mean.

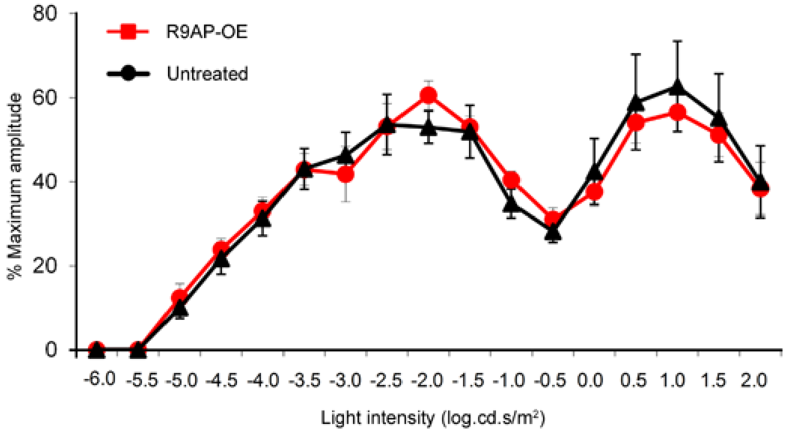


**Figure S3. No effect on 6 Hz ERG intensity-response curve following over-expression of R9AP in wild-type mice**

The eyes treated with rAAV2/8.Rho.mR9ap showed no shift in 6 Hz ERG intensity-response curve compared to that for the untreated contralateral eyes in C57BL6 mice (N = 5). The data is presented as average ± standard error of the mean.
